# Mechanism of Sanliangsan Lipid-Lowering in treating hyperlipidemia: insights from network pharmacology and molecular docking

**DOI:** 10.1101/2025.09.25.678651

**Authors:** Ze Yu, YaMin Hu, Yaqi Wang, Yufei Yang, Wang Lv

## Abstract

**Background and Objective:** This study aimed to identify the active ingredients, key targets, and signaling pathways of Sanliangsan Lipid-Lowering (SLL) in the treatment of hyperlipidemia (HLP) using network pharmacology, and to further clarify the material basis and mechanism responsible for its lipid-lowering efficacy.

**Methods:** All active ingredients of SLL were retrieved from the Traditional Chinese Medicine Systems Pharmacology Database and Analysis Platform (TCMSP), followed by the identification of target proteins corresponding to each active ingredient. Target genes associated with these target proteins were obtained from the Uniprot database, with duplicate genes excluded to generate the predicted targets of the active ingredients.Cytoscape was utilized to construct the “Herb-Active Ingredient-Predicted Target” network for SLL. Targets related to hyperlipidemia (HLP) were retrieved from the GeneCards, OMIM, and DrugBank databases using “hyperlipidemia” as the keyword. The intersection of the predicted targets of SLL active ingredients and HLP-related targets was defined as the potential therapeutic targets of SLL for HLP. Utilizing Cytoscape 3.9.1 software, the top 20 targets with the highest Degree values were selected to construct the “Herb-Active Ingredient-Target-Disease” network, (esignated as the “Herb-Active Ingredient-Target Network of SLL against HLP.” The STRING database was employed to build a protein-protein interaction (PPI) network of the potential therapeutic targets, which was subsequently analyzed using the Network Analysis, CytoNCA, and CytoHubba plugins in Cytoscape to identify the top 10 key targets. The Metascape platform was used for Gene Ontology (GO) functional analysis and Kyoto Encyclopedia of Genes and Genomes (KEGG) pathway enrichment analysis of the key targets, and the Bioinformatics online tool was applied to visualize the analytical results. Furthermore, AutoDock Tools 1.5 software was utilized to perform molecular docking between the active ingredients and key targets, thereby validating the results of the network pharmacology analysis.

**Results:** A total of 115 active ingredients and 255 corresponding predicted targets of SLL were screened from the TCMSP database. Additionally, 2106 HLP-related targets were retrieved from the aforementioned databases, and 140 common targets (potential therapeutic targets) were identified through intersection analysis. From these common targets, the top 20 with the highest Degree values were selected for the construction of the “Active Ingredient-Potential Target-Disease” network. Network analysis revealed that kaempferol, quercetin, and formononetin were the top three active ingredients in terms of content, while tumor necrosis factor (TNF), interleukin-6 (IL6), protein kinase B1 (AKT1), interleukin-1β (IL1B), prostaglandin-endoperoxide synthase 2 (PTGS2), peroxisome proliferator-activated receptor γ (PPARG), caspase 3 (CASP3), and hypoxia-inducible factor 1α (HIF1A) were the targets associated with the largest number of active ingredients. The PPI network consisted of 140 nodes and 3,207 edges. By using the CytoHubba plugin for analysis, six key targets were identified: AKT1, TNF-α, IL-1 β, IL6, PPARG and PTGS2. The GO functional enrichment analysis revealed 30 entries, which encompassed 10 biological processes (such as positive regulation of transcription by RNA polymerase II), 10 cellular components (such as extracellular space), and 10 molecular functions (such as transcription coactivator binding). Additionally, 10 KEGG signaling pathways were identified, including Lipid and atherosclerosis, the AGE-RAGE signaling pathway in diabetic complications, and Fluid shear stress and atherosclerosis. The findings indicated that SLL exerts a therapeutic effect on HLP by regulating multiple signaling pathways. Molecular docking results demonstrated that the binding energies of the three core ingredients—kaempferol, quercetin, and formononetin—with all key targets were less than -5.0 kJ/mol, with the lowest binding energy (-10.1 kJ/mol) observed between formononetin and PTGS2, indicating a strong binding affinity between these two molecules.

**Conclusion:** This study identified the potential active ingredients, key targets, and core signaling pathways of SLL in treating HLP, confirming that SLL exerts its lipid-lowering effect through a synergistic “multi-component, multi-target, multi-pathway” mechanism. It provides a theoretical basis for further experimental validation and clinical application of SLL in managing HLP.

## Introduction

Hyperlipidemia (HLP) is a prevalent metabolic disorder among middle-aged and elderly populations, with an increasing incidence rate. It also serves as a major risk factor for atherosclerotic cardiovascular and cerebrovascular diseases^[1]^. Although statins and fibrates demonstrate significant lipid-lowering effects, long-term administration is associated with safety concerns such as rhabdomyolysis and abnormal liver function ^[2]^. Sanliangsan Lipid-Lowering (SLL), a traditional Chinese medicine (TCM) folk empirical prescription, is composed of *Astragalus membranaceus* (Huang-Qi)*, Hawthorn* (Shan-Zha), *Angelica sinensis* (Dang-Gui), *Glycyrrhiza uralensis* (Gan-Cao), *Lonicera japonica* (Jin-Yin-Hua) and *Citrus aurantium* (Zhi-Shi) ^[3]^. Clinical and animal studies have confirmed its remarkable lipid-regulating efficacy ^[4–9]^ . However, its “multi-component, multi-target, multi-pathway” molecular mechanism is rarely reported. In this study, network pharmacology analysis combined with molecular docking validation was employed to explore the molecular mechanism underlying the therapeutic effect of SLL on HLP, aiming to provide theoretical support and reference for the clinical application of SLL in HLP treatment and subsequent experimental research (Figure 1).

**Figure 1.**
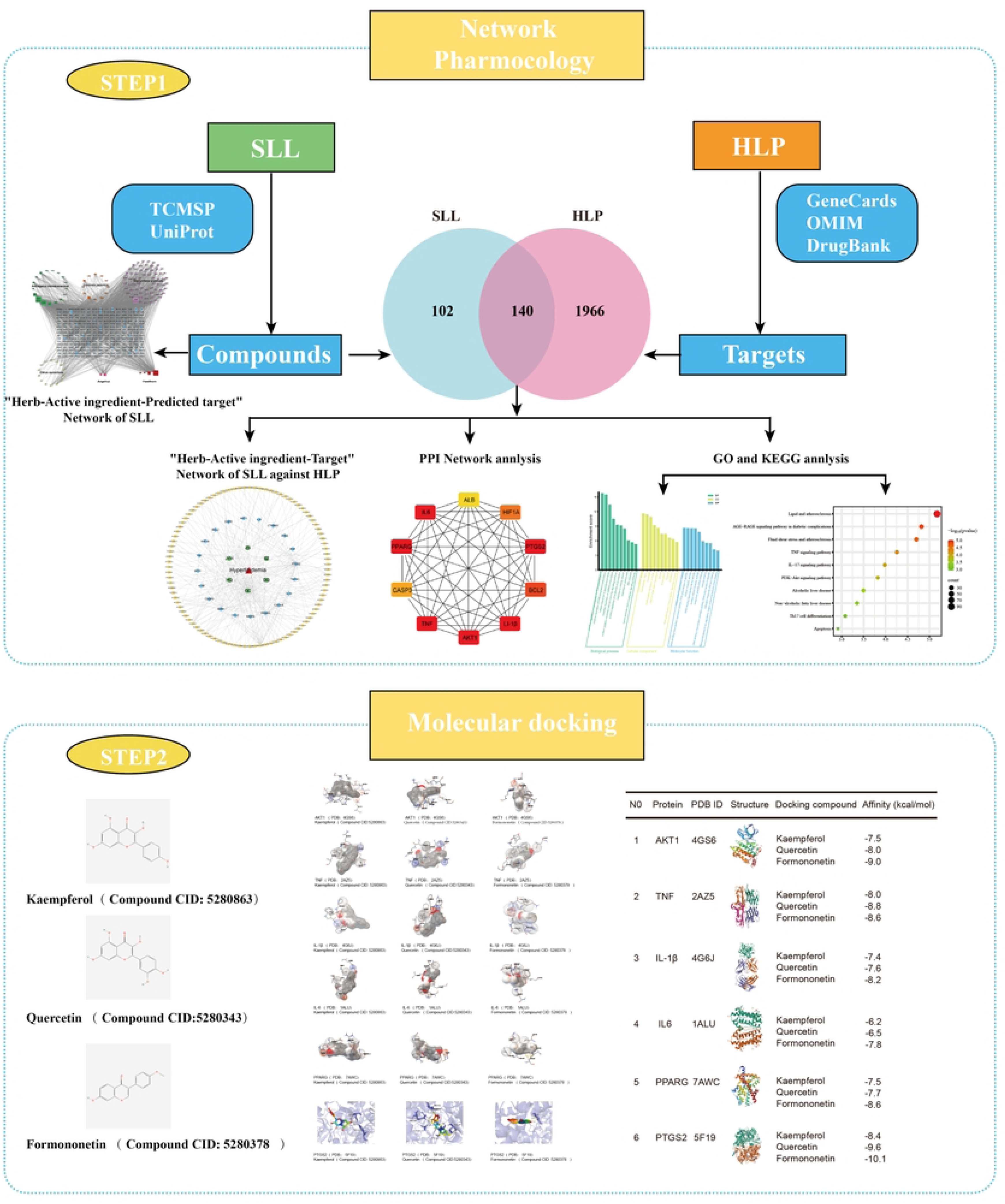
Research Process.

## 1 Materials and Methods

### 1.1 Screening of Active Ingredients in SLL and Prediction Targets

All components of SLL, including those derived from *Astragalus membranaceus* (Huang-Qi)*, Hawthorn* (Shan-Zha), *Angelica sinensis* (Dang-Gui), *Glycyrrhiza uralensis* (Gan-Cao), *Lonicera japonica* (Jin-Yin-Hua), and *Citrus aurantium* (Zhi-Shi), were obtained from the Traditional Chinese Medicine Systems Pharmacology Database and Analysis Platform (TCMSP). Active ingredients in SLL were screened based on the criteria of oral bioavailability (OB) ≥ 30% and drug-likeness (DL) ≥ 0.18^[10]^. The TCMSP database (https://tcmspw.com/tcmsp.php) was used to search for the target proteins corresponding to each active ingredient. Subsequently, the Uniprot database (with the species set to *Homo sapiens*) was utilized to obtain the gene IDs associated with each target protein. Duplicate genes were removed, and the remaining gene IDs were integrated to generate the predicted targets of the active ingredients (Table 1).

### 1.2 Mining of HLP and SLL Related Targets

Genes associated with hyperlipidemia were retrieved from the GeneCards database, (http://www.genecards.org) the OMIM database(http://omim.org), and the DrugBank database (http://www.drugbank.com) using “hyperlipidemia” as the keyword. Targets with a “Score” higher than the median were selected as HLP-related disease targets. The intersection of the predicted targets of SLL active ingredients and HLP-related disease targets was defined as the potential therapeutic targets of SLL for HLP.Using Cytoscape 3.9.1 software, the top 20 targets with the highest degree values were utilized to construct the “Active Ingredient-Potential Target-Disease” network for SLL in the context of HLP treatment.

### 1.3 Protein-Protein Interaction (PPI) Analysis of Potential Therapeutic Targets

The STRING database (https://cn.string-db.org) was used to calculate the PPI of the potential therapeutic targets of SLL for HLP. A PPI network was constructed using these potential targets, with the species restricted to Homo sapiens and a confidence threshold set to > 0.4. The PPI results were imported into Cytoscape 3.9.1 software, and the Network Analysis plugin was utilized to conduct a topological analysis of the PPI network. The median values of degree, closeness, and betweenness were used as screening criteria ^[11]^. The CytoNCA plugin was employed to rank the targets within the PPI network, and the CytoHubba plugin was applied for topological analysis to identify the top 10 key gene targets in the PPI network.

### 1.4 GO Functional and KEGG Pathway Enrichment Analyses of Key Targets

To further investigate the mechanism of action of the key targets of SLL in HLP treatment, the information of the screened key targets was imported into the Metascape platform for GO functional annotation and KEGG pathway enrichment analysis. The results were visualized using the Bioinformatics online tool.(https://www.bioinformatics.com.cn/).

### 1.5 Molecular Docking

Molecular docking was conducted to verify the key targets and core active ingredients identified through the PPI network, assess the binding interactions between targets and ingredients, and preliminarily confirm the outcomes of the network pharmacology analysis.The 3D structures of the core ingredients, available in SDF format, were retrieved from the PubChem database and subsequently converted to PDB format using the Open Babel GUI. AutoDock Tools version 1.5.6 were utilized to determine the minimum number of atoms required for rotation and active bonds, as well as to generate files in the PDBQT format. The protein structures of the key targets were retrieved and downloaded from the PDB database (https://www.rcsb.org). After removing heteroatoms using Notepad software, the protein structures were imported into AutoDock Tools 1.5.6 for charge calculation, hydrogen atom addition, and non-polar ion fusion, and finally exported as files in PDBQT format. The PDBQT files of the core ingredients and key targets were imported into AutoDock Tools 1.5.6 to define the active pocket and set the Gridbox coordinates and dimensions. AutoDock Vina scripts were run to perform molecular docking, and PyMOL software was used to visualize the docking results.

## 2 Results

### 2.1 Construction and Analysis of the “Herb-Active Ingredient-Predicted Target” Network

The primary components of SLL were sourced from the TCMSP database, and 115 active ingredients were identified, comprising 17 from *Astragalus membranaceus* (Huang-Qi), 12 from *Lonicera japonica* (Jin-Yin-Hua), 66 from *Glycyrrhiza uralensis*(Gan-Cao), 14 from *Citrus aurantium* (Zhi-Shi), 2 from *Angelica sinensis*(Dang-Gui), and 4 from *Hawthorn*(Shan-Zha), corresponding to 255 targets.

These active ingredients and their predicted targets were imported into Cytoscape 3.9.1 software to construct the “Herb-Active Ingredient-Predicted Target” network (Figure 2). Among these targets, PTGS2, prostaglandin-endoperoxide synthase 1 (PTGS1), and peroxisome proliferator-activated receptor (PPAR) exhibited the highest Degree values, indicating that they may be significant targets for the lipid-regulating effect of SLL.

**Figure 2.**
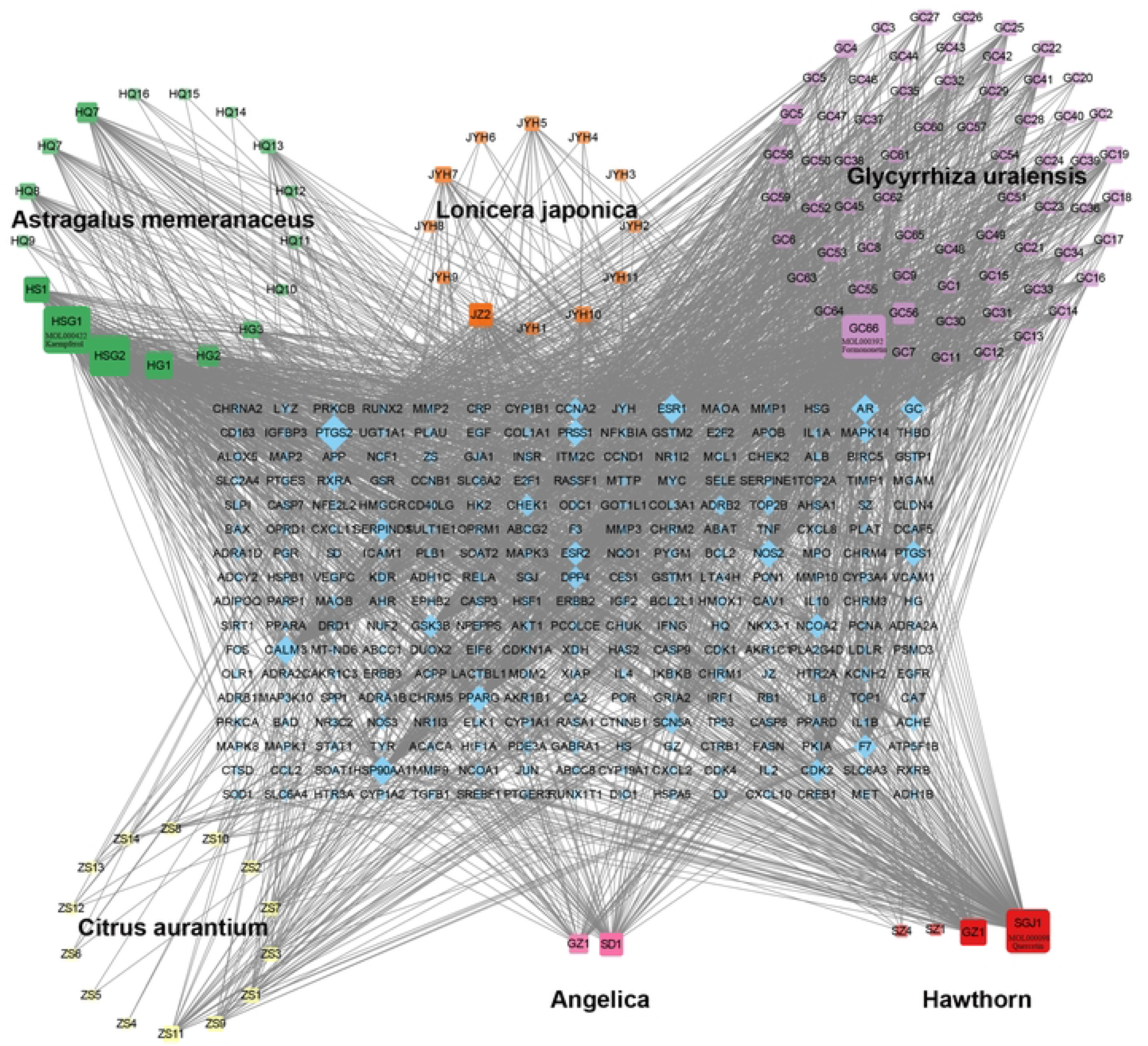
“Herb-Active Ingredient-Predicted Target” Network of SLL. In the figure, square nodes represent active pharmaceutical ingredients: green nodes indicate ingredients derived from *Astragalus membranaceus*(Huang-Qi), *Lonicera japonica*(Jin-Yin-Hua), *Glycyrrhiza uralensis*(Gan-Cao), yellow nodes from *Citrus aurantium* (Zhi-Shi), pink nodes from *Angelica sinensis*(Dang-Gui), and red nodes from *Hawthorn*(Shan-Zha). Diamond nodes with a grid pattern represent disease targets. Lines indicate the targeting relationships between these nodes. The Degree value represents the number of edges connected to a node; a higher Degree value of a predicted target node signifies a stronger association with the active ingredients in SLL.

### 2.2 ”Herb-Active ingredient-Target” Network of SLL against HLP

A total of 2,106 HLP-related targets were screened from the GeneCards, OMIM, and DrugBank databases. Targets with a “Score” higher than the median were selected as HLP-related disease targets. An intersection analysis of the predicted targets of SLL active ingredients and HLP-related targets yielded 140 common targets, which were identified as the potential therapeutic targets of SLL for HLP (Figure 3).The active ingredients and potential therapeutic targets were imported into the Cytoscape 3.9.1 software, and the top 20 targets with the highest degree values were utilized to construct the “Active Ingredient-Potential Target-Disease” network for SLL in the context of HLP treatment.The network comprised 122 nodes, which included 95 compounds, 20 target genes, 6 herb nodes, and 1 disease node. The Analyze-Network function was utilized to calculate the network degrees.It was determined that kaempferol, quercetin, and formononetin were the top three active ingredients in terms of content. Furthermore, TNF was the gene associated with the largest number of active ingredients, succeeded by IL6, AKT1, IL1B, PTGS2, PPARG, CASP3, and HIF1A.(Figure 4, Table 2).

**Figure 3.**
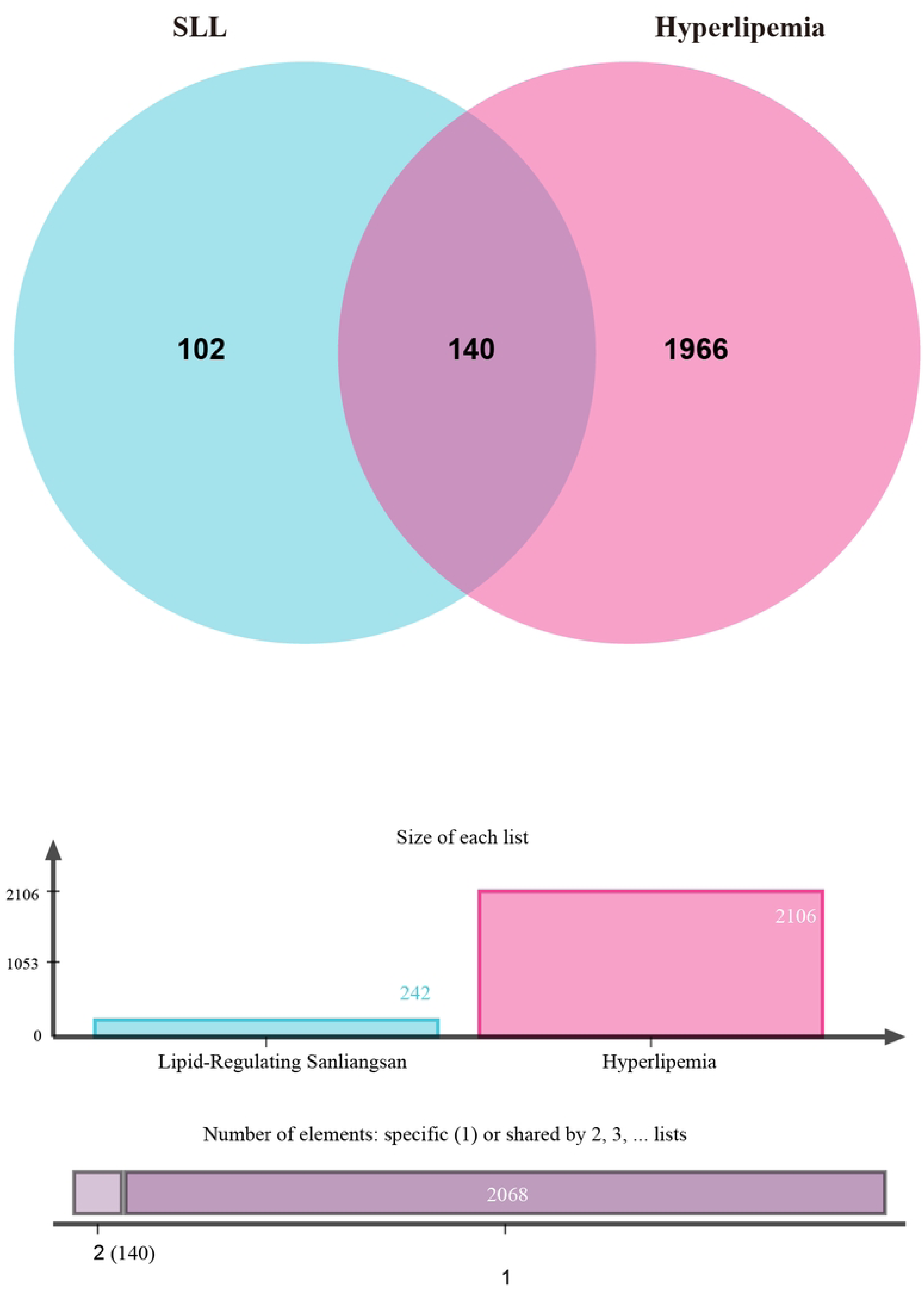
Potential Therapeutic Targets of SLL for HLP. The intersection of the predicted targets of SLL active ingredients (shown in blue) and HLP-related disease targets (shown in red) yielded 140 common targets, which are the potential therapeutic targets of SLL for HLP.

**Figure 4.**
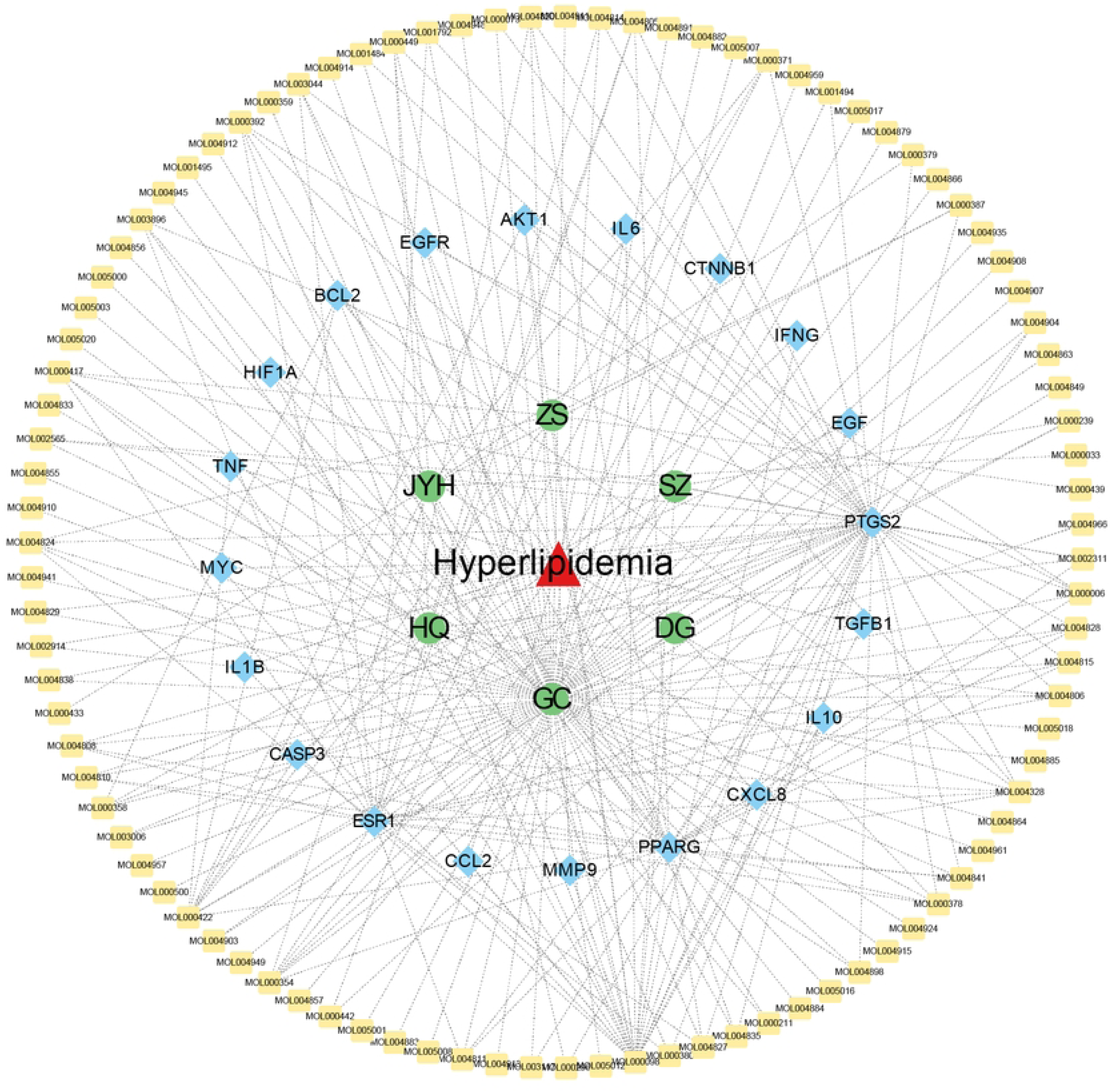
“Active Ingredient-Potential Target-Disease” Network of SLL against HLP. In the figure, red triangular nodes represent the disease (HLP), green circular nodes represent SLL herbs, blue diamond nodes represent potential therapeutic targets, and peripheral yellow square nodes represent active ingredients. Edges denote the relationships between active ingredients, potential therapeutic targets, and the disease; the connections between nodes represent their functional relationships. A higher number of connections to a node signifies a more significant role of the target or compound within the network.

### 2.3 Network analysis and core target screening of PPI

The PPI network for SLL’s HLP treatment comprised 140 nodes and 3207 edges (Figure 5A). The CytoNCA plugin was utilized to rank the targets within the PPI network, maintaining the same number of nodes and edges (Figure 5B).The CytoHubba plugin was utilized to reconstruct the network’s core, consisting of 10 nodes and 46 edges. Ultimately, a subnetwork was obtained, centered around six key targets: AKT1, TNF-α, IL-1β, IL6, PPARG, and PTGS2 (Figure 5C).

**Figure 5.**
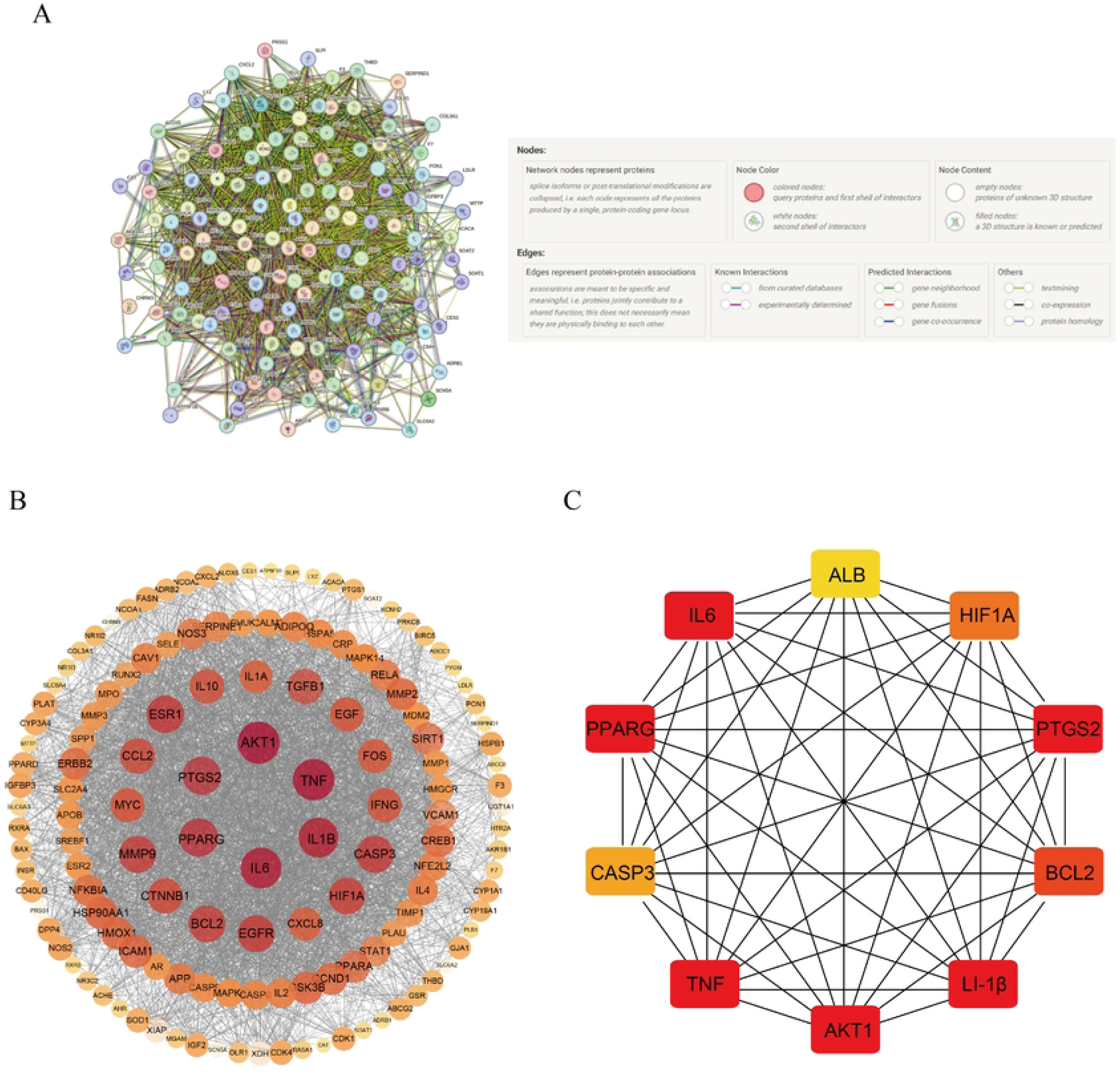
Protein-Protein Interaction (PPI) Network and Key Subnetwork. (A) Protein-Protein Interaction (PPI) network;(B) Network derived from CytoNCA analysis;(C) Key subnetwork of the top 10 nodes identified by CytoHubba.

### 2.4 GO Functional and KEGG Pathway Enrichment Analyses

The Metascape platform was utilized to conduct gene enrichment analysis on the 140 core nodes, encompassing biological processes (BP) in GO (e.g., positive regulation of transcription by RNA polymerase II), cellular components (CC) (e.g., extracellular space), and molecular functions (MF) (e.g., transcription coactivator binding) (Figure 6A). The KEGG pathway enrichment analysis revealed 71 KEGG pathways, and the top 10 pathways were chosen based on P-values (with the x-axis representing the number of genes within each pathway). The KEGG pathways were primarily enriched in Lipid metabolism and atherosclerosis (involving 91 genes), the AGE-RAGE signaling pathway in diabetic complications (involving 27 genes), and Fluid shear stress and atherosclerosis (involving 27 genes) (Figure 6B).

**Figure 6.**
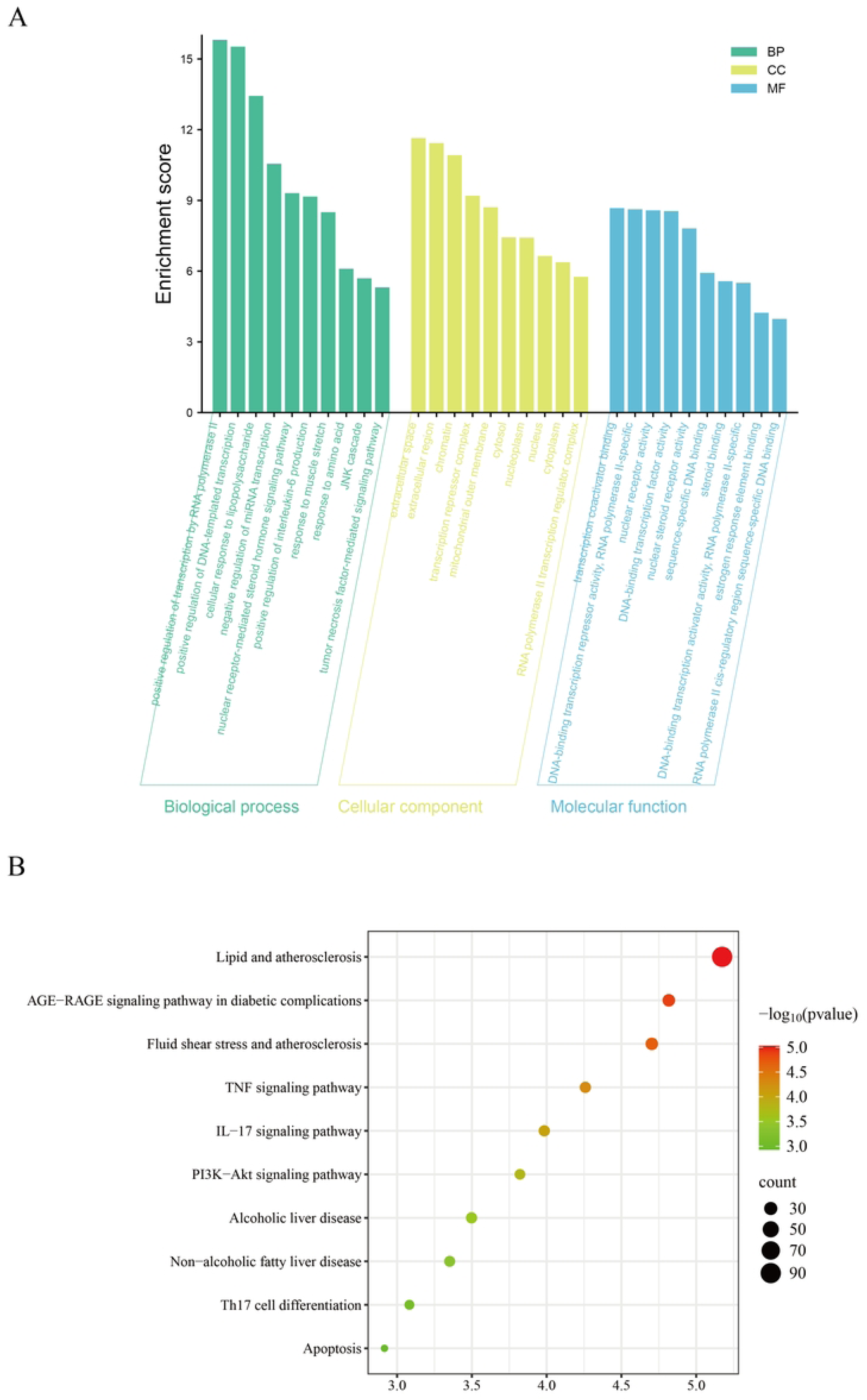
GO and KEGG enrichment analysis. (A) GO enrichment analysis of the target genes. Biological Process (BP, green bars), Cellular Component (CC, yellow bars), and Molecular Function (MF, blue bars). The x-axis represents different enrichment terms, and the y-axis denotes the enrichment score. (B) KEGG Enrichment Analysis. In the KEGG pathway bubble chart, the color of the bubbles shifts from red to green, indicating an increase in Log10(P) values; the size of the bubbles corresponds to the number of genes enriched within each pathway.

### 2.5 Molecular docking

Based on the results presented in Sections 2.3 and 2.4, molecular docking was conducted between six key targets (AKT1, TNF-α, IL-1β, IL-6, PPARG, and PTGS2) and three core ingredients (kaempferol, quercetin, and formononetin). The binding ability of these molecules was evaluated using binding energy values. The results indicated that the binding energies of all compounds with the target proteins were less than -5.0 kJ/mol, signifying a high binding capacity between these compounds and target proteins (Figure 7A). Among these pairs, the binding energy between formononetin and PTGS2 was -10.1 kJ/mol, representing the optimal binding combination (Figure 7B).

**Figure 7.**
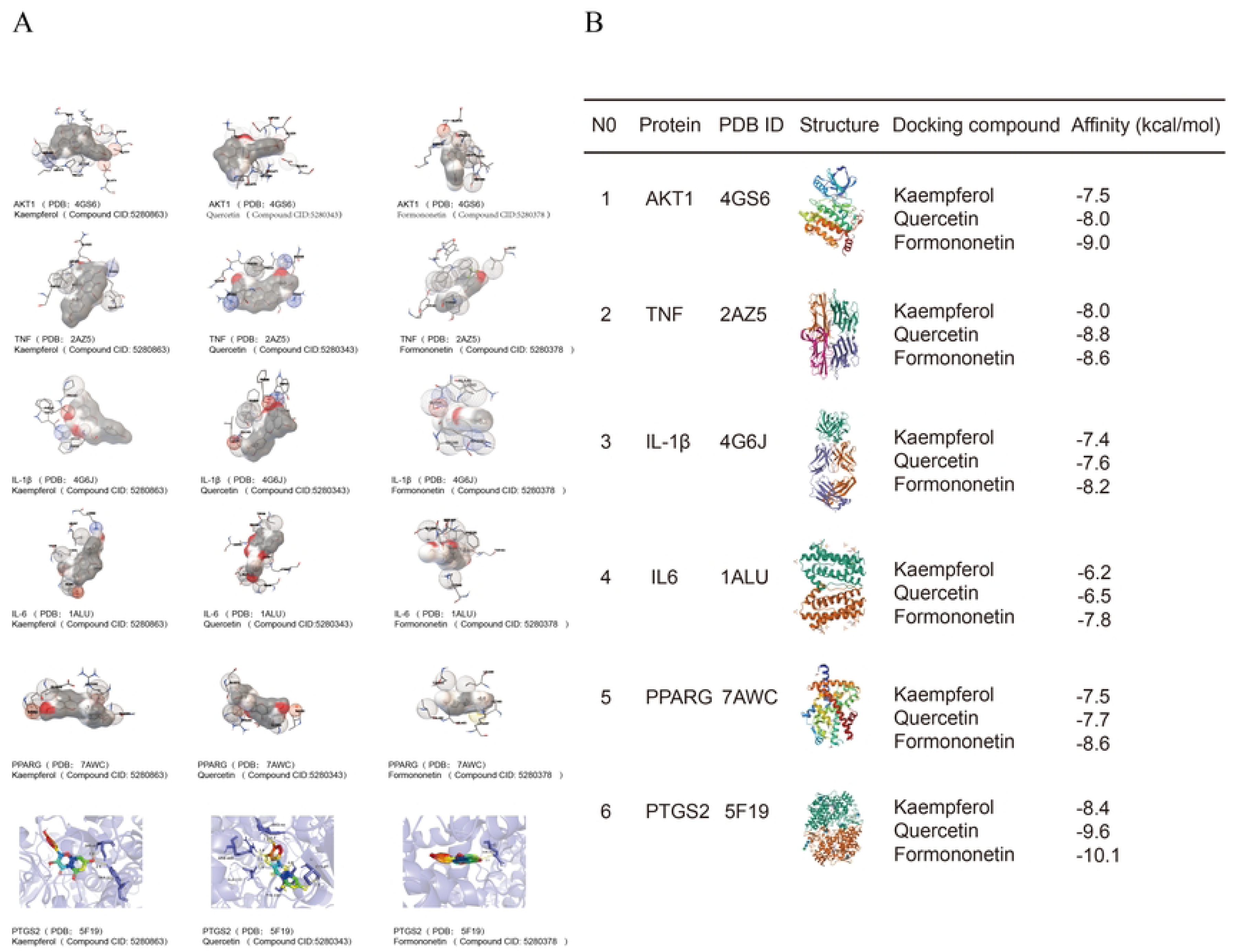
Molecular Docking of SLL’s Key Targets in HLP. (A) The 3D binding modes of kaempferol, quercetin, and formononetin with their target proteins (AKT1, TNF-α, IL-1β, IL-6, PPARG, and PTGS2); (B) A summary of the molecular docking affinities (expressed in kcal/mol) between kaempferol, quercetin, formononetin and each target protein (AKT1, TNF-α, IL-1β, IL-6, PPARG, and PTGS2), including the corresponding protein PDB IDs and structural diagrams.

## Discussion

This study revealed that as a compound traditional Chinese medicine (TCM) preparation, the efficacy of SLL in treatin HLP is attributed to the synergistic effect of its multiple components. Kaempferol, quercetin, and formononetin were identified as the core active ingredients of SLL for treating HLP. All three belong to the flavonoid class, which has been reported to regulate lipid metabolism and exert anti-atherosclerotic effects^[12–14]^ .

Kaempferol can alleviate oleic acid-induced lipid accumulation and oxidative stress in HepG2 cells ^[15]^ , and exert anti-inflammatory effects by inhibiting the nuclear factor-κB (NF-κB) pathway as well as suppressing the expression of inflammatory factors such as interleukin-6 (IL-6), interleukin-1β (IL-1β), and tumor necrosis factor-α (TNF-α) ^[16–17]^ .Quercetin can activate peroxisome proliferator-activated receptor γ (PPARγ) in the liver, thereby promoting the transport of cholesterol from macrophages to high-density lipoprotein (HDL) and reducing foam cell formation ^[18]^ . Additionally, quercetin can scavenge reactive oxygen species (ROS) and inhibit the activation of NF-κB ^[19]^ , which mitigates HLP-related oxidative stress and chronic inflammation, further delaying the progression of atherosclerosis ^[20]^ . A recent study indicated that formononetin may be effective in treating atherosclerosis by regulating the Kruppel-like factor 4 (KLF4)-sterol regulatory element-binding protein RNA (SREBP RNA) interaction to attenuate its development ^[21]^ .

This study demonstrated that SLL, with its diverse active ingredients, acts on multiple key targets involved in the pathogenesis of HLP, namely AKT1, TNF-α, IL-1β, IL-6, PPARG, and PTGS2. Among these targets, AKT1 can inhibit the activation of sterol regulatory element-binding protein 1c (SREBP-1c) in the liver, thereby reducing triglyceride synthesis^[22]^ .Conversely, the inhibition of AKT1 activity leads to the upregulation of pro-inflammatory mediators, such as IL-6, promotes the polarization of macrophages towards an inflammatory phenotype, and accelerates the progression of atherosclerosis ^[23]^ . TNF-α serves as a critical target in HLP-related chronic inflammation; it inhibits the transcription of the lipoprotein lipase (LPL) gene by disrupting the binding of nuclear factor Y (NF-Y) to octamer-binding proteins. This disruption reduces LPL synthesis, inhibits triglyceride hydrolysis, and results in lipid accumulation in adipocytes, thereby further exacerbating HLP ^[24–25]^ .As a core member of the pro-inflammatory cytokine family, IL-1β is closely associated with hepatic steatosis and vascular injury induced by HLP ^[26]^ . Elevated levels of IL-6 are linked to dyslipidemia and an increased risk of atherosclerosis; as a downstream product of inflammatory factors such as TNF-α and IL-1β, IL-6 can further amplify inflammatory signals through autocrine and paracrine mechanisms ^[27]^ . PPARG, a member of the nuclear receptor superfamily, transcriptionally activates the expression of lipolytic genes (e.g., adipose triglyceride lipase [ATGL] and hormone-sensitive lipase [HSL]), thereby enhancing lipolysis in adipose tissue and counteracting obesity and metabolic disorders ^[28]^ .It also inhibits the expression of inflammatory factors, such as TNF-α and IL-6, in macrophages, reduces foam cell formation, and delays atherosclerosis progression ^[29]^ . Prostaglandin-endoperoxide synthase 2 (PTGS2) plays a key role not only in inflammatory signal-related pathological processes but also affects intra-plaque cellular homeostasis and lipid deposition, making it a potential intervention target for atherosclerosis induced by HLP ^[30–31]^ .

This study further identified that SLL exerts its therapeutic effect on HLP by regulating multiple pathways, including lipid metabolism and atherosclerosis, the AGE-RAGE signaling pathway in diabetic complications, fluid shear stress and atherosclerosis, the PI3K-Akt signaling pathway, the IL-17 signaling pathway, the TNF signaling pathway, Th17 cell differentiation, and apoptosis.Relevant studies have indicated that the lipid and atherosclerosis pathway facilitates the transport of cholesterol from macrophages to HDL, diminishes cholesterol accumulation in vascular walls, and hinders the activation of the NF-κB signaling pathway. Consequently, this process suppresses foam cell formation and delays the onset and progression of atherosclerotic plaques ^[32–33]^ .The activation of the AGE-RAGE signaling pathway is a key mechanism underlying vascular injury in patients with HLP. HLP facilitates the accumulation of advanced glycation end products (AGEs) in vascular endothelial cells, which subsequently activates the NF-κB and ROS production pathways and upregulates the expression of inflammatory factors, such as TNF-α and IL-6 ^[34–35]^ . The Fluid shear stress pathway is closely linked to HLP, as it affects the development and progression of atherosclerosis by modulating endothelial cell gene expression and phenotypic transitions ^[36–37]^ . The activation of the TNF signaling pathway can induce oxidative stress, enhance the transcytosis of low-density lipoproteins (LDLs) in endothelial cells, and foster the development of atherosclerosis ^[38]^ . This study also indicates that the primary compound targets of SLL are concentrated in liver-related pathways, including Alcoholic liver disease and Non-alcoholic fatty liver disease. This suggests that SLL may have a synergistic effect on regulating lipid synthesis, lipid breakdown, inflammatory responses, and vascular function through multiple pathways, such as Lipid and atherosclerosis, AGE-RAGE, and Fluid shear stress and atherosclerosis, to ameliorate HLP. Additionally, SLL may exert potential regulatory effects on cellular physiological processes and liver diseases.

In conclusion, this study utilized network pharmacology and molecular docking to preliminarily elucidate the synergistic “multi-component, multi-target, multi-pathway” mechanism of SLL in treating HLP. The core active ingredients—kaempferol, quercetin, and formononetin—can bind to key targets (such as AKT1, TNF-α, IL-1β, IL-6, PPARG, and PTGS2, and subsequently regulate multiple pathways, including those related to lipid and atherosclerosis, AGE-RAGE, and fluid shear stress and atherosclerosis. This regulation synergistically modulates lipid metabolism, inflammatory responses, and vascular function, thereby improving HLP. This study provides a reference for further experimental and clinical research on SLL in the treatment of HLP; however, the findings still require further biological validation.

